# Genomic Analysis of Human Noroviruses Using Hybrid Illumina-Nanopore Data

**DOI:** 10.1101/2021.03.30.437798

**Authors:** Annika Flint, Spencer Reaume, Jennifer Harlow, Emily Hoover, Kelly Weedmark, Neda Nasheri

**Affiliations:** Genomics Laboratory, Bureau of Microbial Hazards, Health Canada, Ottawa, ON, Canada; National Food Virology Reference Centre, Bureau of Microbial Hazards, Health Canada, Ottawa, ON, Canada; Department of Biochemistry, Microbiology and Immunology, Faculty of Medicine, University of Ottawa, ON, Canada

**Keywords:** Norovirus, full-length amplicons, Illumina MiSeq, Oxford Nanopore, single nucleotide variant, recombination

## Abstract

Whole genome sequence (WGS) analysis of noroviruses is routinely performed by employing a metagenomic approach. While this methodology has several advantages, such as allowing for examination of co-infection, it has some limitations such as the requirement of high viral load to achieve full-length or near full-length genomic sequences. In this study, we used an amplification approach to obtain full-length genomic amplicons from 39 Canadian GII isolates followed by deep sequencing on Illumina and Oxford Nanopore platforms. This approach significantly reduced the required viral titre to obtain full-genome coverage. Herein, we compared the coverage and sequences obtained by both platforms and provided an in-depth genomic analysis of the obtained sequences, including the presence of single nucleotide variants (SNVs) and recombination events.

## Introduction

Noroviruses are the most common agent causing acute gastroenteritis leading to an estimated 684 million illnesses and approximately $60 billion in societal costs worldwide (Bartsch, et al. 2016). There are currently no vaccines or therapeutics licensed against norovirus (Cates, et al. 2020). Norovirus transmission primarily occurs by person-to-person contact through the fecal-oral route or contaminated food and surfaces (Teunis, et al. 2015).

Noroviruses belong to the *Caliciviridae* family and have a 7.5 kb, positive sense, single-stranded RNA genome that is enclosed in a non-enveloped icosahedral capsid (Green. 2013). The genome is organized into three open reading frames (ORFs). ORF1 encodes a polyprotein that is cleaved into six non-structural viral proteins, including the RNA-dependent RNA polymerase (RdRp). ORF2 encodes VP1, the major structural capsid protein, and ORF3 encodes VP2, a minor structural capsid protein (Green. 2013). Noroviruses are considered fast-evolving viruses (de Graaf, et al. 2016), and their genomes are tremendously diverse due to accumulation of point mutations and recombination (Parra. 2019).To date, noroviruses are classified into at least 10 genogroups. Noroviruses are further classified into at least 49 genotypes based on the diversity of ORF2 and 60 types based on the diversity of the RNA-dependent-RNA polymerase (RdRP) gene (Chhabra, et al. 2019).

Whole genome sequencing (WGS) of noroviruses, which is often carried out through metagenomic approaches, has allowed for source attribution, lineage analysis, identification of recombination events and variant analysis (Parra and Green. 2015, Nasheri, et al. 2017, Petronella, et al. 2018). However, high viral titre is required to obtain full-genome coverage through metagenomics approaches (Nasheri, et al. 2017). Alternatively, amplicon-based sequencing using sequence-specific primer sets decreases the viral load requirement in the sample, but introduces amplification bias and assumes conserved viral synteny, which could lead to overlooking genomic variations (Cotten, et al. 2014). Parra and colleagues have recently developed a method to amplify full-length norovirus GII genomes followed by the use of the Illumina platform to obtain deep sequencing data on the full-genome amplicons (Parra, et al. 2017).

Third generation sequencing devices such as Oxford Nanopore’s MinION, which can produce long reads up to 100’s of kilobases, has become the method of choice to elucidate viral recombination and to identify subgenomic sequences (Viehweger, et al. 2019). On the other hand, second-generation sequencing technologies like Illumina, despite a low error rate, are restricted by read-length (≤300 nt) which, even using paired-end 2×300 sequencing, considerably complicates the investigation of recombination and identification of subgenomic sequences. However, adoption of MinION sequencing for routine surveillance of viruses has been limited due to concerns of sequence accuracy (Bull, et al. 2020). To overcome this concern, we performed amplicon-based long-read MinION and short-read paired-end Illumina MiSeq on matched norovirus-positive stool samples.

The aim of this study is to combine Illumina and Oxford Nanopore sequencing to reconstruct a highly accurate consensus sequence of the norovirus isolates and to provide insight into the viral recombination and genetic diversity.

## Materials and Methods

### Samples

The norovirus GII positive samples are from the British Columbia Centre for Disease Control (BMH19-089 to BMH19-137), obtained in winter and spring 2019, and from the archive of the National Food Virology Reference Centre at Health Canada (BMH11-021, BMH12-030, BMH 13-039, BMH14-054, BMH14-056, BMH15-059, BMH15-063, BMH16-074, BMH16-077, BMH18-086, and BMH18-087), which were obtained from winter 2011 to spring 2018. The presence of norovirus GII RNA was confirmed by droplet digital PCR (Bio-Rad, Hercules, California, USA) using the probes and primers that were described previously (Nasheri, et al. 2017, Petronella, et al. 2018). The samples either were from an outbreak or were sporadic. An outbreak includes >2 epidemiologically linked cases with >1 norovirus-positive sample. This study has been granted an exemption from requiring ethics approval by Health Canada and a formal consent was not required because the study participants were anonymized.

### RNA Extraction and Full-length Amplicon Generation

Full-length amplicons were generated as described before (Parra, et al. 2017). Briefly, 10% stool suspensions were clarified by centrifugation (6000 × g for 5 min) and the supernatant was filtered through a 0.45 μM then 0.22 μM filter (Millipore, Etobicoke, Ontario, Canada). RNA was extracted from filtrate using the MagMax Viral RNA Isolation Kit (Ambion) following manufacturer’s recommendations. Complementary DNA was synthesized from the viral RNA using the Tx30SXN primer (GACTAGTTCTAGATCGCGAGCGGCCGCCCTTTTTTTTTTTTTTTTTTTTTTTTTTTTTT (Katayama, et al. 2002), and the Maxima H Minus First Strand cDNA Synthesis Kit (Thermo Fisher Scientific). Amplification of the full-length genome was performed using a set of primers that target the conserved regions of the 5’-and 3’-end of GII noroviruses (GII1-35: GTGAATGAAGATGGCGTCTAACGACGCTTCCGCTG, and Tx30SXN), and the SequalPrep Long PCR Kit (Invitrogen) following manufacturer’s recommendations.

### Viral RNA Load Determination

Viral titres were determined by droplet digital PCR (Bio-Rad, Hercules, California, USA) using the probes and primers that were described previously (Nasheri, et al. 2017, Petronella, et al. 2018).

### Sanger Sequencing

Samples that did not generate full-length replicons after 3 attempts were subjected to dual typing by Sanger sequencing. For this purpose, we performed conventional RT-PCR targeting of a 570 bp product that includes the 3’-end of the polymerase region and 5’-end of the major capsid gene followed by Sanger sequencing, as described previously (Cannon, et al. 2019).

### Illumina Sequencing

Norovirus libraries were constructed using the NexteraXT DNA Library Preparation Kit and Nextera XT Index Kit v2 according to the manufacturer’s instructions (Illumina Inc.). Paired-end Illumina sequencing was performed on a MiSeq instrument (v3 chemistry, 2 × 300 bp) according to manufacturer instructions (Illumina Inc.).

### Oxford Nanopore Sequencing

Norovirus cDNA samples were treated with 0.16 mg/mL RNase A (100 mg/mL, Qiagen) for 30 minutes at 37 °C. Samples were subsequently size selected using modified SPRI magnetic beads to remove DNA fragments < 1500 bp (Hosomichi, et al. 2014). Briefly, 1mL of Ampure XP beads (Beckman Coulter) was applied to a magnetic stand and the supernatant was discarded. The beads were resuspended in 0.5 mL of 20% PEG, 2.5 M NaCl solution. The modified SPRI beads were added to DNA samples at a 0.35× volume and incubated for 5 minutes at room temperature. Samples were applied to a magnetic stand, the supernatant discarded and beads washed 2× with 125 μL 80% ethanol. The beads were air dried for 30 seconds followed by incubation for 2 minutes at room temperature in 45 μL H_2_O. Samples were applied to a magnetic stand and the eluted DNA was quantified using a Qubit 4 fluorometer (Fisher).

DNA libraries were constructed using the Ligation Sequencing Kit 1D (SQK-LSK108) and the PCR Barcoding Expansion 1-12 Kit (EXP-PBC001) according to the manufacturer protocol (Oxford Nanopore Technologies, Oxford Science Park, UK). Twelve barcoded libraries were pooled per run and sequenced on a 1D MinION device (R9.4, FLO-MIN106) using FLO-MIN106 flow cells for up to 24 hrs. Signal processing, base calling, demultiplexing and adapter trimming were performed using the Guppy (Guppy GPU v 3.3.3+fa743ab).

### Bioinformatic Analysis

#### Read processing, de novo whole genome assembly and genome annotation

Illumina dataset quality was assessed by FastQC (v0.11.8) followed by read processing using Fastp (v 0.20.0) (Chen, et al. 2018) to remove adapter and barcode sequences, correct mismatched bases in overlaps, and filter low-quality reads. Nanopore dataset quality was analyzed using NanoPlot (v1.20.0) (De Coster, et al. 2018) and full genome length Nanopore reads 7.5 -7.7 kb were extracted using NanoFilt (v2.7.1) (De Coster, et al. 2018). Nanopore reads were taxonomically classified using Kaiju (v1.7.3) (Menzel, et al. 2016) and the viral Kaiju database (NCBI RefSeq database curated 25/05/2020). The top scoring read was extracted using Seqtk (v1.3, https://github.com/lh3/seqtk) and non-norovirus GII specific reads were discarded.

Error correction of the full genome length Nanopore read for each sample was performed using the consensus function of Medaka (v1.1.3, https://github.com/nanoporetech/medaka) and Medaka model r941_min_fast_g303 to polish the sequence using Nanopore long reads. The sequence was further polished using Pilon (v1.23) (Walker, et al. 2014) and Illumina short reads using default parameters to obtain the consensus sequence. The whole genome sequences were genotyped using the Norovirus Genotyping Tool v2.0 (Kroneman, et al. 2011) and annotated using Vapid (v1.6.6) (Shean, et al. 2019) and the Vapid viral database (NCBI complete viral genomes curated 01/05/2018).

#### Coverage plots

Illumina and Nanopore reads were mapped to the norovirus *de novo* whole genomes using BBMap (v38.18, https://sourceforge.net/projects/bbmap). Depth of coverage was assessed using the Samtools depth function (v1.7, https://github.com/samtools/samtools), and data was graphed using GraphPad (v6.01).

#### Phylogenetic analysis

Nucleotide sequences for ORF1, ORF2 and ORF3 from each norovirus strain and closely related GenBank reference sequences (MK753033, KJ407074, MT409884, MH218671, KC576912, MK762635, KX158279, MT731279, MN996298, KU757046) were aligned with MUSCLE using MEGA (v10.1.8) (Tamura, et al. 2013). Maximum Likelihood phylogenetic trees based on the Tamura-Nei model were constructed and visualized in MEGA using the aligned sequences and 1000 bootstrap replicates.

#### Single nucleotide variant (SNV) analysis

Single nucleotide variants were identified for each norovirus sample with Breseq (v0.35.5) (Deatherage and Barrick. 2014) using the mutation prediction pipeline and default parameters. Variants were identified using the norovirus Illumina reads for each sample and highly similar complete genome GenBank references. GenBank accessions used were MK753033, KJ407074, MT409884, MH218671, KC576912, MK762635, KX158279, MT731279, MN996298, and KU757046.

#### Recombination analysis

Whole genome assemblies for the 39 isolates and closely related GenBank reference sequences (MK753033, KJ407074, MT409884, MH218671, KC576912, MK762635, KX158279, MT731279, MN996298, KU757046) were aligned with MUSCLE using MEGA (v10.1.8) (Tamura, et al. 2013). Recombination breakpoints and identification of potential parental sequences were performed using the Recombination Detection Program (RDP4) (v4.101) (Martin, et al. 2015) using seven recombination methodologies: RDP, GENECONV, MaxChi, Bootscan, Chimera, SiScan and 3Seq. A sliding window of 200-bp and a step size of 20 bp, and a multiple-comparison corrected *p* < 0.05 were used. Graphs were created using Simplot within RDP4.

#### Data availability

The complete *de novo* genome sequences of the 39 norovirus isolates used in this study have been uploaded to GenBank under accession numbers: MW661246 to MW661284. All SRAs are available in GenBank under BioProject ID PRJNA713985.

## Results

### Amplicon Production

To examine whether full-length amplicons could be generated for a variety of GII samples, we employed the primers that encompass 5’UTR to 3’UT on 57 GII positive samples (44 isolated in 2019 and the remaining 13 were archived samples isolated since 2011). Full-length amplicons were obtained from 39 samples, and multiple efforts for the remaining 18 samples failed to generate any full-length amplicons. Thus, we conducted conventional RT-PCR to obtain partial amplicons for dual typing of these samples by Sanger sequencing. Four out of 18 samples failed to generate any RT-PCR product for dual typing. The full-length amplicons obtained from all 39 samples were subjected to both Illumina and Nanopore sequencing as depicted in Figure 1. The data obtained from both approaches were used to assemble the full-genomes and for further sequence analysis.

**Figure 1.**
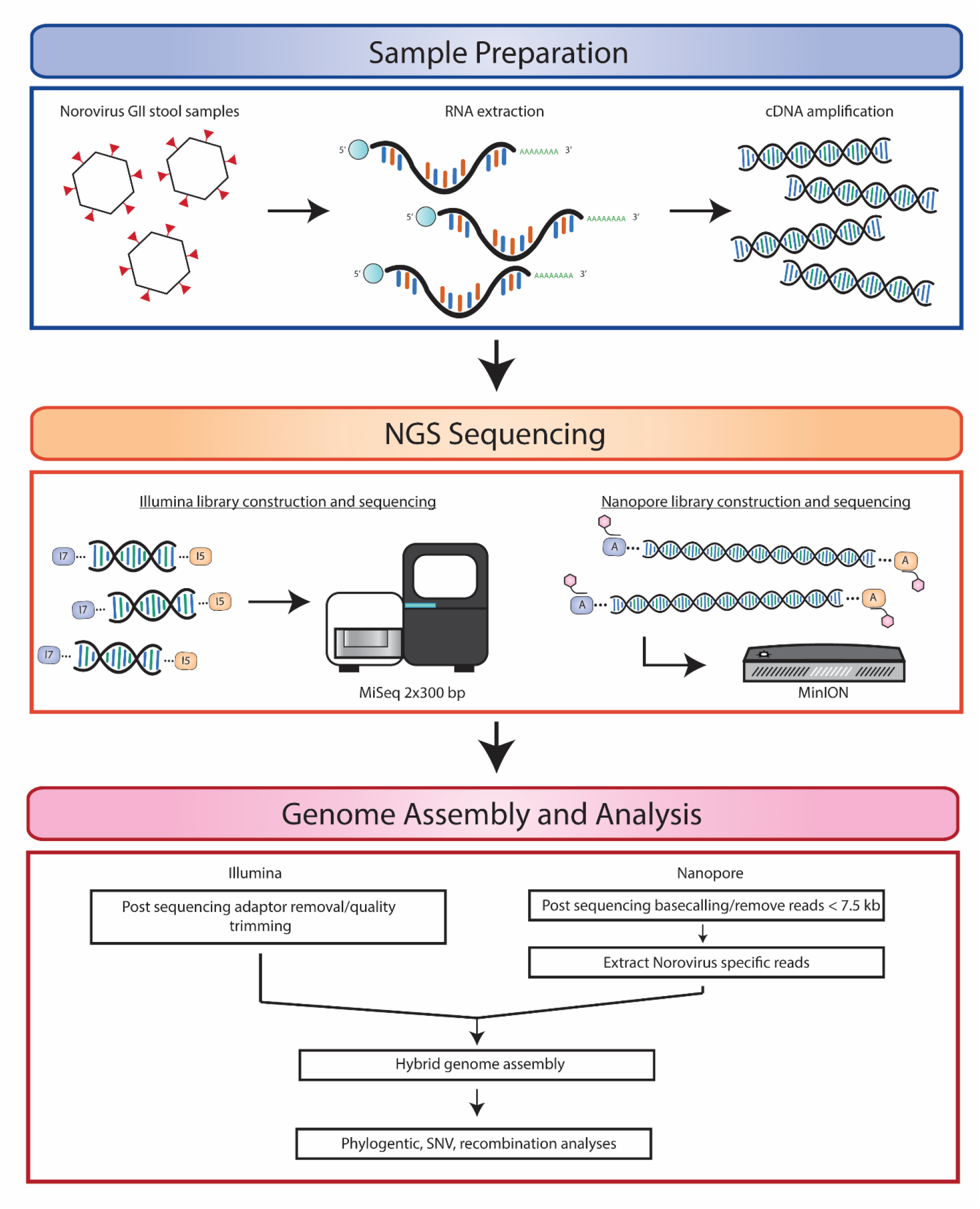
Schematic representation of experimental approach for norovirus *de novo* whole genome sequencing, assembly and bioinformatics analysis. Norovirus viral RNA was extracted from stool samples followed by full-length cDNA amplification. cDNA was sequenced on an Illumina MiSeq platform and an Oxford Nanopore MinION device to obtain high quality short read and long read data, respectively. Illumina and Nanopore reads were processed to remove adaptor and barcode sequences followed by quality trimming of Illumina reads and length filtering of Nanopore reads. A hybrid approach using both Illumina and Nanopore reads was used to produce *de novo* full-length norovirus genomes. Genome annotation and downstream analyses were performed using the norovirus assemblies.

As shown in Figure 2 and Supplementary Table 1, GII.P16 (63%) was the dominant polymerase type followed by GII.P12 (11%), GII.P7 (10%), and GII.P31 (8%). As expected, GII.4 (44%) was the most prevalent genotype, followed by GII.1 (17%), GII.3 (13%), GII.2 (12%), and GII.6 (8%). The dominant GII.P16 was mostly associated with GII.4, making GII.4 [P16] the most prevalent strain (30%), however, it was also associated with GII.1, and GII.2 (Supplementary Table 1).

**Figure 2.**
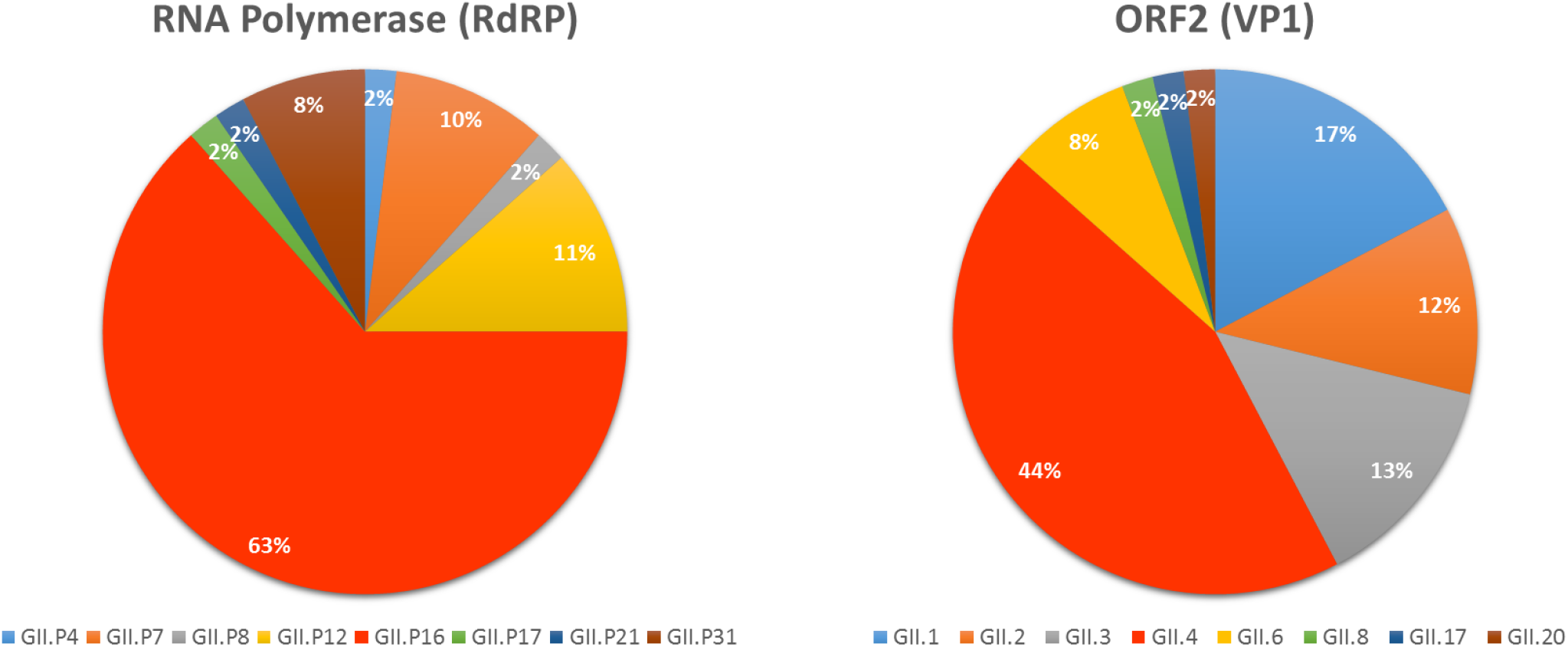
Norovirus genotypes identified in this study. Norovirus Genotyping Tool v2.0 was used to determine the polymerase type (A) and the capsid/ORF2 genotype (B) of all the samples sequenced (by Sanger or next-generation sequencing) in this study.

### Analysis of the Assay Sensitivity

To determine the lowest viral genome copy number that would produce full-length amplicons, we made serial dilutions for four representative samples; BMH16-77, BMH18-86, BMH19-95, and BMH19-96. The full-length amplicons generated from the highest dilution (lowest viral load) was subjected to Illumina sequencing to ensure the full-genome sequence could still be obtained. As shown in Table 1, the lowest viral RNA titre that generated full-length sequence ranged from 1.7 to 3.4 ×10^2^ RNA copy number.

**Table 1.** The lowest RNA copy number that generated full-length amplicons

| Sample ID | RNA titre | Lowest titre generated amplicon |
| --- | --- | --- |
| BMH16-77 | 4.90E+06 | 2.70E+02 |
| BMH18-86 | 2.20E+06 | 2.10E+02 |
| BMH19-95 | 3.40E+06 | 1.70E+02 |
| BMH19-96 | 3.50E+07 | 3.40E+02 |

### Comparison between Illumina and MinION Coverage

To compare sequencing coverage along the full length of the norovirus genomes, representative samples from each genotype identified in this study were selected for coverage depth analysis. For each genotype, the Illumina and Nanopore reads were mapped to their corresponding consensus sequence and the sequencing depth at each nucleotide position was determined. As shown in Figure 3, the Illumina and Nanopore read data produced similar patterns of coverage depth across the length of the norovirus genomes. For the representative genotypes GII.1[P16], GII.4[P4], GII.4[P16] and GII.4[P31], the Nanopore data showed increased sequencing depth across ~2500 bp of ORF1 of the genome. A similar trend is also observed in the Illumina sequencing data, although not as pronounced. The Illumina and Nanopore data for GII.2[P16] showed increased depth of coverage across the first ~2500 bp of ORF1 and the last ~2500 bp corresponding to ORF2 and ORF3. Interestingly, the Nanopore data for GII.8[P8] had a large increase in depth across ~500 bp at the 5’ end of the genome in contrast to the Illumina data, which showed consistent sequencing depth across the entire length of the genome (excluding the 5’ and 3’ ends of the genome). Consistent sequencing depth coverage across the genome was observed for GII.3[P12], GII.3[P21], GII.6[P7], and GII.17[P17] for both sequencing technologies.

**Figure 3.**
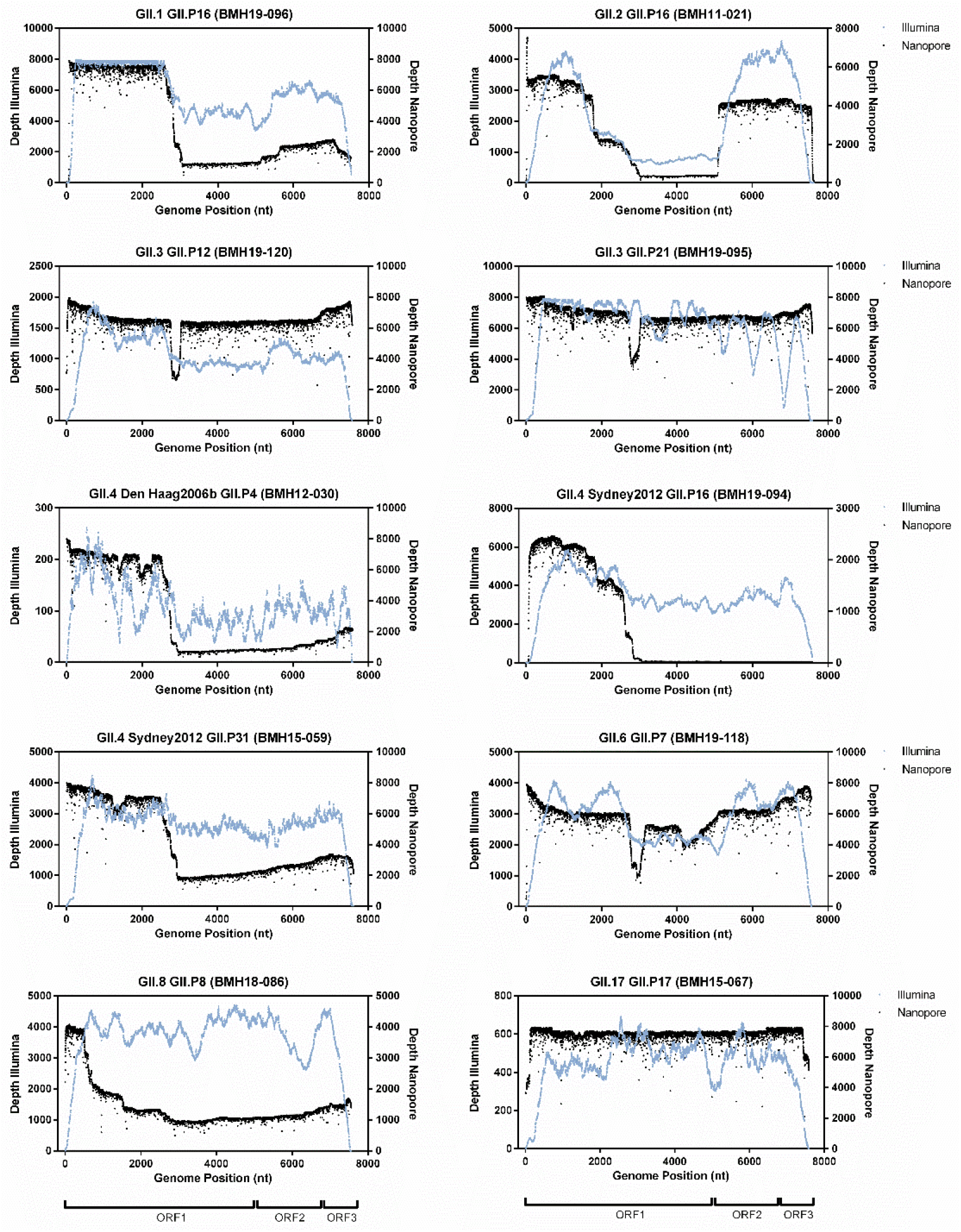
Coverage profiles of representative norovirus genotypes using Illumina and Nanopore read data. Reads were mapped to the *de novo* consensus sequence for each genotype and read type. Depth across the genome at each nucleotide position for Illumina (blue) and Nanopore (black) is shown. Schematic representation of the norovirus genome for each ORF is illustrated below each column of graphs.

Overall, the Illumina data typically lacked coverage at both the 5’ and 3’ end of the norovirus genomes (Figure 3) resulting in incomplete genome assemblies and only partial coding sequence data for ORF1 (data not shown). In contrast, the Nanopore reads yielded full-length sequences, which covered both the 5’ and 3’ ends of the norovirus genome. Additionally, using the Nanopore reads it was possible to obtain complete and/or partially complete sequences of the 5’ and 3’ UTR regions.

### Phylogenetic Analysis

Phylogenetic analysis of the 39 Canadian *de novo* assembled norovirus strains from this study was performed for ORF1, ORF2, and ORF3 (Figure 4A). The phylogenetic tree for ORF1 is based on the nucleotide sequence of the 6 non-structural genes p48, NTPase, p22, VPg, Pro, and RdRP. Each ORF from the 39 norovirus strains and highly similar reference strains for each genotype were aligned, and Maximum Likelihood trees were constructed. For ORF1, the 39 norovirus samples belong to 8 different polymerase types (GII.P4, GII.P7, GII.P8, GII.P12, GII.P16, GII.P17, GII.P21, GII.P31; Supplementary Table 1). As shown in Figure 4, the majority of the samples (24/39) belong to polymerase type GII.P16 and cluster together (BMH11-021, BMH14-054, BMH16-074, BMH18-085, BMH19-090, BMH19-092, BMH19-093, BMH19-094, BMH19-096, BMH19-097, BMH19-108, BMH19-109, BMH19-110, BMH19-111, BMH19-112, BMH19-113, BMH19-115, BMH19-117, BMH19-125, BMH19-127, BMH19-128, BMH19-129, BMH19-132, BMH19-137). These strains also cluster closely with the NCBI reference strains for the GII.P16 genotype (GenBank accession MK753033, KJ407074, and MK762635). Furthermore, strains BMH19-108, BMH19-109, BMH19-110, BMH19-111, BMH19-112, and BMH19-113 are highly similar and isolated from the same outbreak. Strains BMH13-039, BMH15-059, BMH15-063 and BMH19-130, which belong to GII.P31, clustered together along with the reference strain (GenBank accession KX158279). The GII.P12 sequences (BMH16-077, BMH19-100 and BMH19-120) and reference (GenBank accession MT409884) clustered together. The GII.P7 strains (BMH14-056, BMH19-118, BMH19-119, BMH19-145) and reference strain (GenBank accession MT731279) clustered, although strains BMH14-056 and BMH19-145 show less similarity to strains BMH19-118, BMH19-119. The remaining strains BMH12-030 (GII.P4), BMH18-086 (GII.P8), BMH15-067 (GII.P17) and BMH19-095 (GII.P21) all clustered with their corresponding reference strains (GenBank accession KC576912, MN996298, KU757046 and MH218671 respectively).

**Figure 4.**
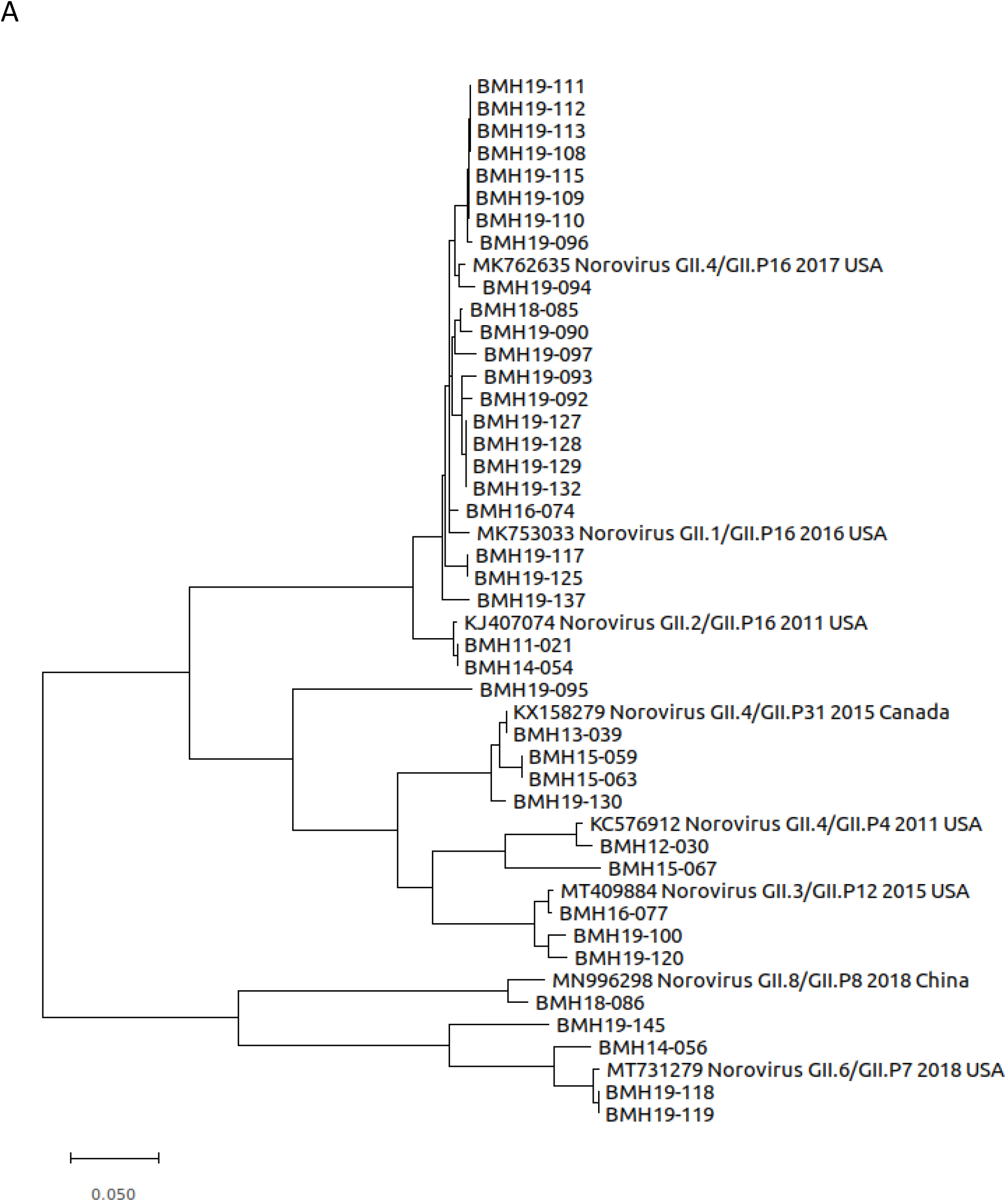
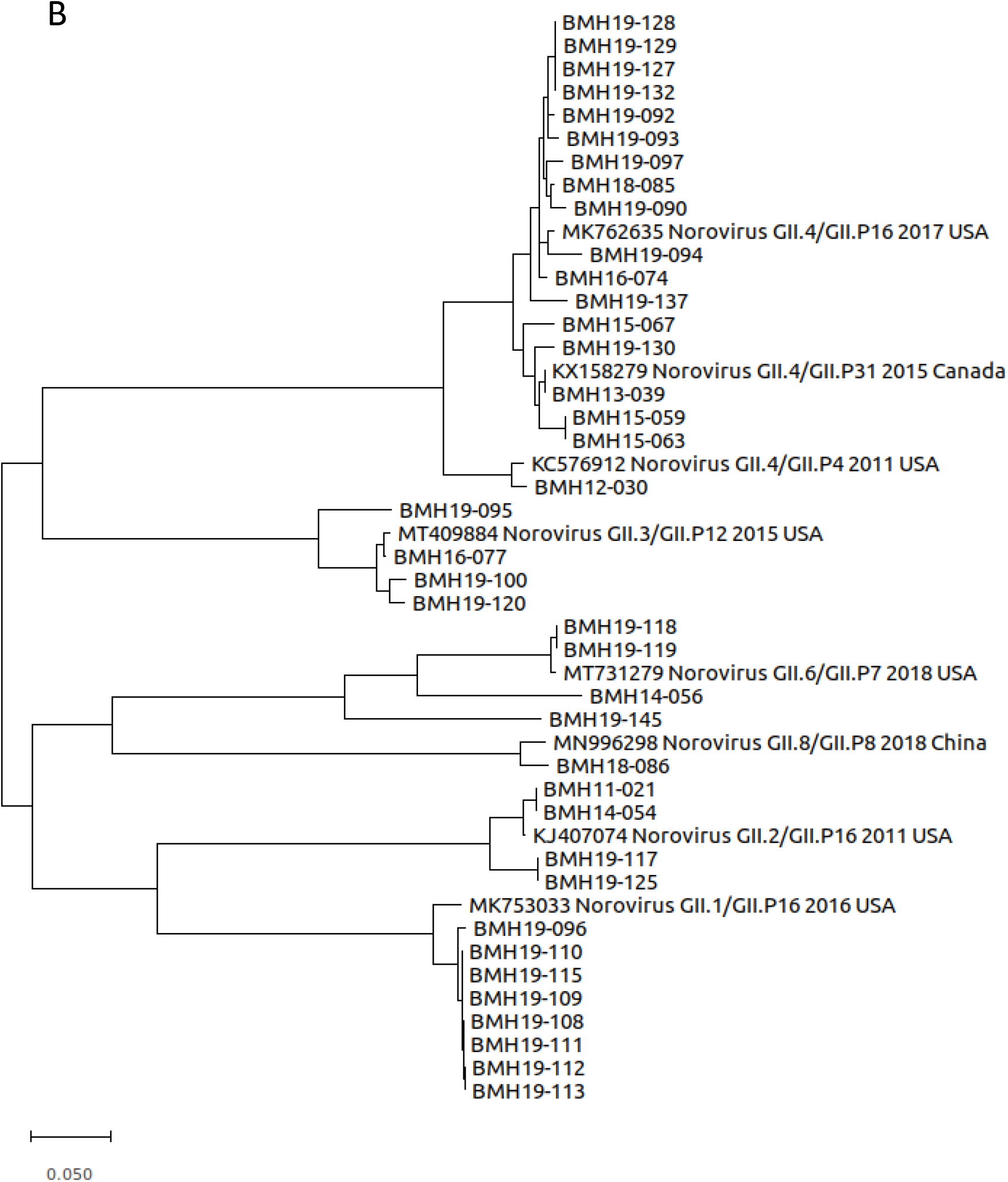
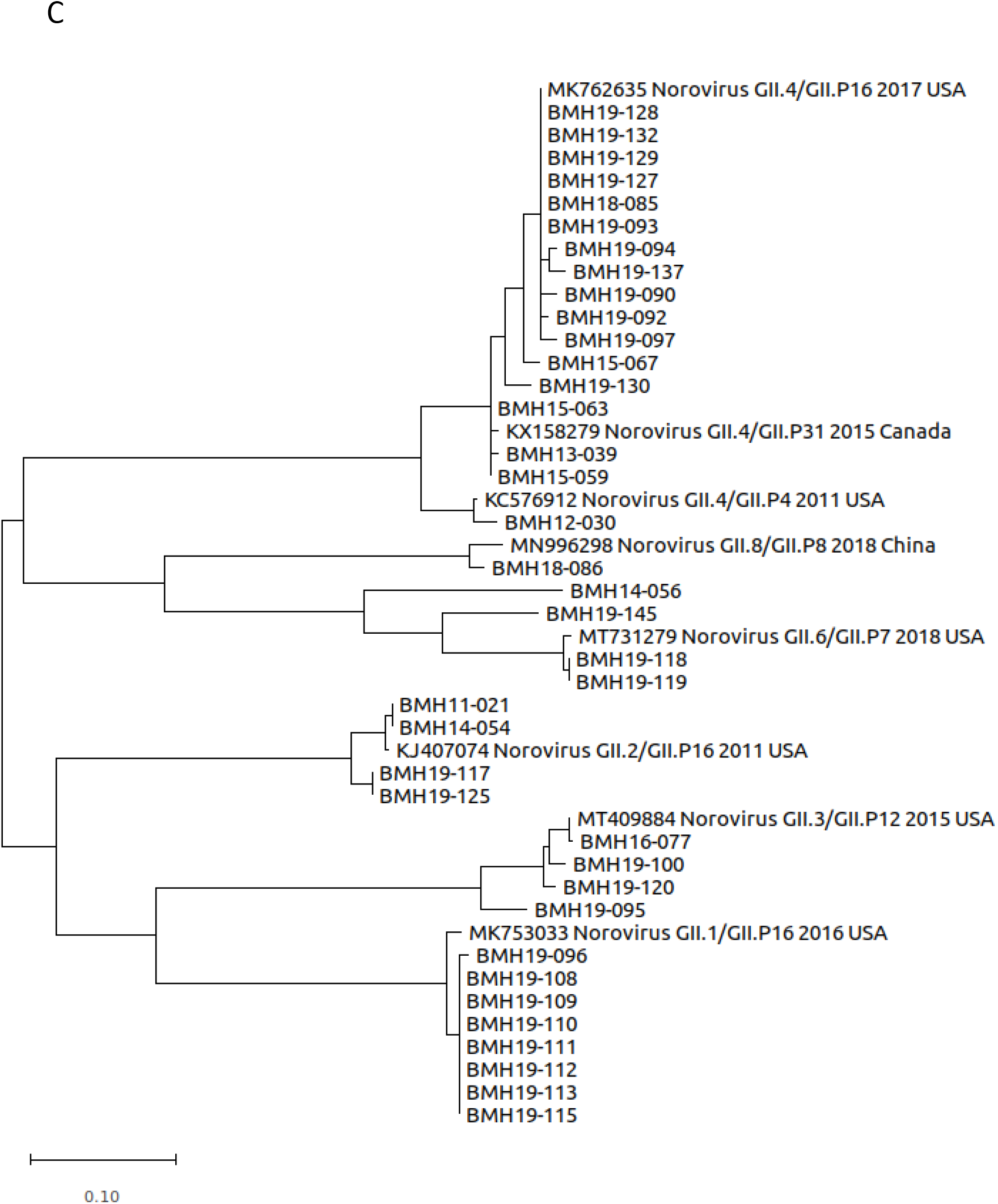
Phylogenetic trees of Canadian norovirus GII ORF1, ORF2 and ORF3 nucleotide sequences obtained from this study. Maximum likelihood trees were constructed using MEGA (v10.1.8) and 1000 bootstrap replicates. The scale bars represent the phylogenetic distance expressed as nucleotide substitutions per site. Reference sequences were obtained from GenBank with accession numbers, genotype, year and country of isolation shown.

The phylogenetic tree for ORF2 (Figure 4B) was constructed using the aligned nucleotide sequences corresponding to the shell, P1 and P2 major capsid domains. The norovirus strains belong to the 7 genotypes GII.1, GII.2, GII.3, GII.4, GII.6, GII.8 and GII.17 with the majority of the strains identified as GII.1 (8/39) and GII.4 (17/39). The GII.1 strains BMH19-096, BMH19-108, BMH19-109, BMH19-110, BMH19-111, BMH19-112, BMH19-113 and BMH19-115 were highly similar where BMH19-108, BMH19-109, BMH19-110, BMH19-111, BMH19-112 and BMH19-113 are all associated with the same norovirus outbreak (reference strain GenBank accession MK753033). The norovirus GII.4 strains, belonging to subtype Sydney 2012, BMH13-039, BMH15-059, BMH15-063, BMH16-074, BMH18-085, BMH19-090, BMH19-092, BMH19-093, BMH19-094, BMH19-097, BMH19-127, BMH19-128, BMH19-129, BMH19-130, BMH19-132, and BMH19-137 also clustered closely together along with the reference strains (GenBank accession MK762635 and KX158279). The remaining GII.4 strain (BMH12-030) belongs to subtype Den Haag 2006b, which shows less similarity to the GII.4 Sydney 2012 strains. The GII.2 norovirus strains (BMH11-021, BMH14-054, BMH19-117 and BMH19-125) clustered together (GenBank accession KJ407074). The ORF2 sequences for BMH16-077, BMH19-095, BMH19-100 and BMH19-120 GII.3 strains clustered with the reference sequence (GenBank accession MT409884), although BMH19-095 showed greater genetic distance from the rest of the strains. For the GII.6 genotype, BMH19-118 and BMH19-119 are highly similar whereas BMH14-056 and BMH19-145 display greater genetic differences (GenBank accession MT731279). The remaining ORF2 genotypes GII.8 (BMH18-086) and GII.17 (BMH15-067) clustered with their respective references (GenBank accession MN996298 and KU757046).

For ORF3, the phylogenetic tree is similar to that observed for ORF2 with all 39 norovirus strains clustering with their respective genotypes GII.1, GII.2, GII.3, GII.4, GII.6, GII.8 and GII.17 (Figure 4C).

### Variant Analysis

The structural domains of the capsid include an N-terminus, a highly conserved shell, and two protruding spike domains (P1 and hypervariable P2) (Smith and Smith. 2019, Debbink, et al. 2012). The capsid is also an antigenic target for the host immune system. Consequently, norovirus mutations to these capsid structural proteins aid in host immune system evasion (Lindesmith, et al. 2013). To investigate potential ORF2 variations in the norovirus samples, single nucleotide variant (SNV) analysis was conducted for each strain using highly similar GenBank references (Figure 5, Supplementary Table 2). From Figure 5, the majority of the SNVs resulting in amino acid changes were observed in the P2 domain. This trend was observed for all of the genotypes in this study. For GII.1, the linked outbreak samples BMH19-108, BMH19-109, BMH19-110, BMH19-111, BMH19-112 and BMH19-113 all have changes in the P2 domain at T325M, H374Q and N293S relative to the reference strain (GenBank accession MK753033). BMH19-117 and BMH19-125, samples from genotype GII.2, showed changes in the N-terminus (S24N), P1 (V256I), P2 (V319I, V335I, V373I), and P1 (V440I) (GenBank accession KJ407074). The majority of samples (17/39) in this study belong to the GII.4[P16] genotype (BMH16-074, BMH18-085, BMH19-090, BMH19-092, BMH19-093, BMH19-094, BMH19-097, BMH19-127, BMH19-128, BMH19-129, BMH19-132, BMH19-137), and with the exception of BMH19-137, contain a common V377A mutation in the P2 domain relative to the reference strain (GenBank accession MK762635). The norovirus GII.6 strains, BMH19-118, BMH19-119 and BMH19-145 all have a S174P change in the conserved shell domain, as well as a G354Q variation in the P2 domain of BMH19-118 and BMH19-119 (GenBank accession MT731279).

**Figure 5.**
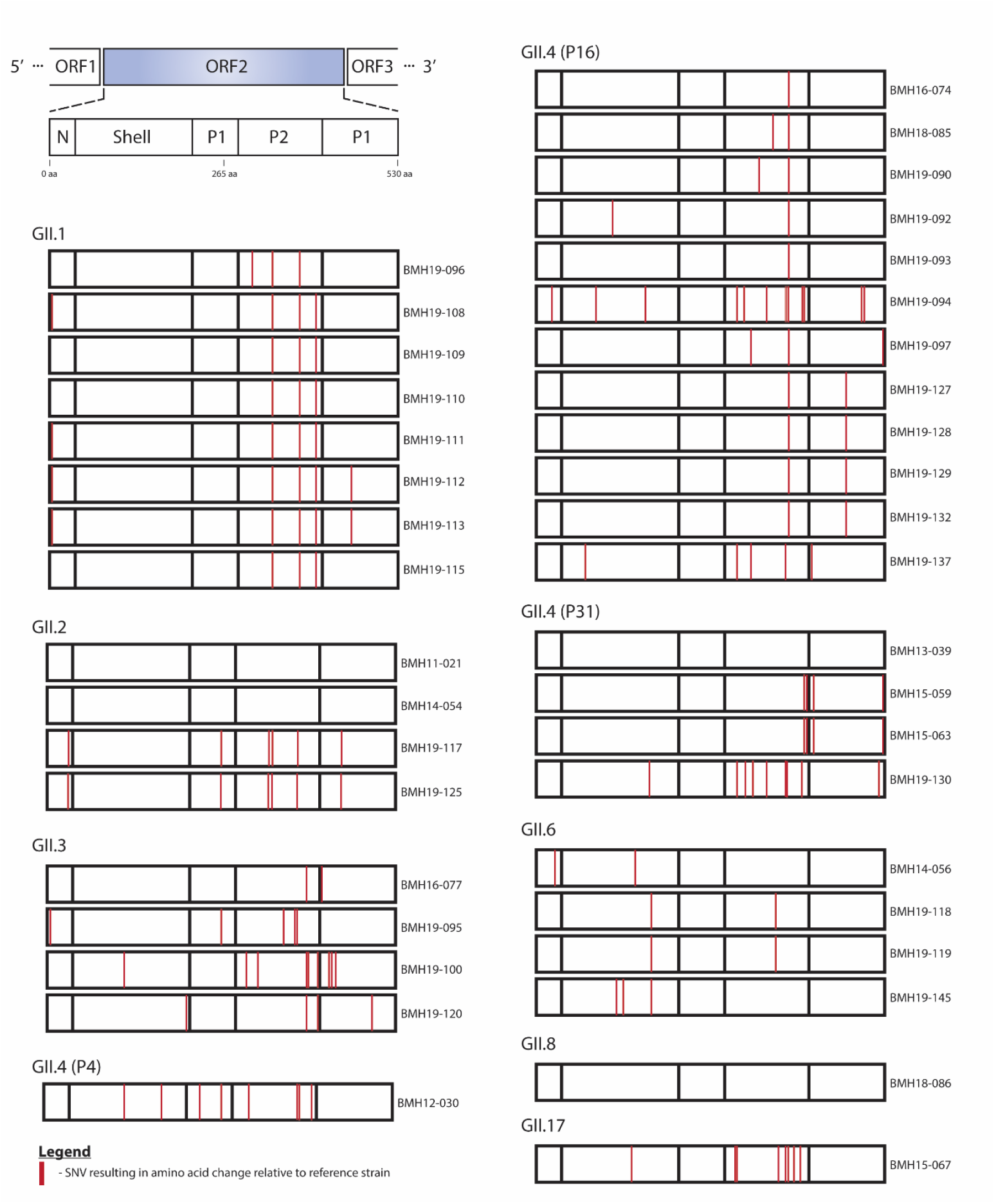
Single nucleotide variation (SNV) analysis of norovirus ORF2 major capsid region. Norovirus *de novo* assemblies were assessed for variants relative to highly similar NCBI reference sequences. Schematic representation of the norovirus ORF2 region is shown (upper left). SNVs that resulted in non-synonymous amino acid changes are shown (red lines) at corresponding amino acid positions in the N-terminal, Shell, P1, and P2 domains for each isolate. Isolates are grouped by capsid genotype. The GenBank reference Isolates (accession numbers in brackets) used were GII.1-2016-USA (MK753033), GII.2-2011-USA (KJ407074), GII.3-2015-USA (MT409884), GII.3-2015-UK (MH218671), GII.4-2011-USA (KC576912), GII.4-2017-USA (MK762635), GII.4-2015-Canada (KX158279), GII.6-2018-USA (MT731279), GII.8-2018-China (MN996298), and GII.17-2016-China (KU757046).

The norovirus RdRp (NS7) is the core enzyme for RNA replication. Its structure is highly similar to those of other positive-strand RNA viruses and can be described as a partially closed right hand, with fingers, thumb, and palm subdomains (Deval, et al. 2017). The fingers and palm domains interact to form a channel. Within this channel there are 7 conserved motifs named A through G, which interact with the template, the nascent RNA, and the NTPs for RNA synthesis and comprise the active site (Deval, et al. 2017). As shown in Figure 6, the majority of the SNVs resulting in amino acid changes in RdRP were observed in the Fingers and the Palm domains. Also, as previously mentioned, GII.P16 was the dominant polymerase type, associated with three capsid genotypes; GII.4, GII.2, and GII.1. Similar to what has been observed for the VP1 protein, the linked outbreak samples BMH19-108, BMH19-109, BMH19-110, BMH19-111, BMH19-112 and BMH19-113 all have the same changes in the Finger and Palm domains at T1294S, S1401T, V1446A relative to the reference strain (GenBank accession MK753033) (Supplementary Table 3). The K1646R substitution is one of the few amino acid changes in the Thumb domain that was observed in the majority of the GII.4[P16] isolates, while the S1401T change in the Palm domain was observed for most GII.P16 isolates regardless of the capsid genotype (Figure 6 and Supplementary Table 3).

**Figure 6.**
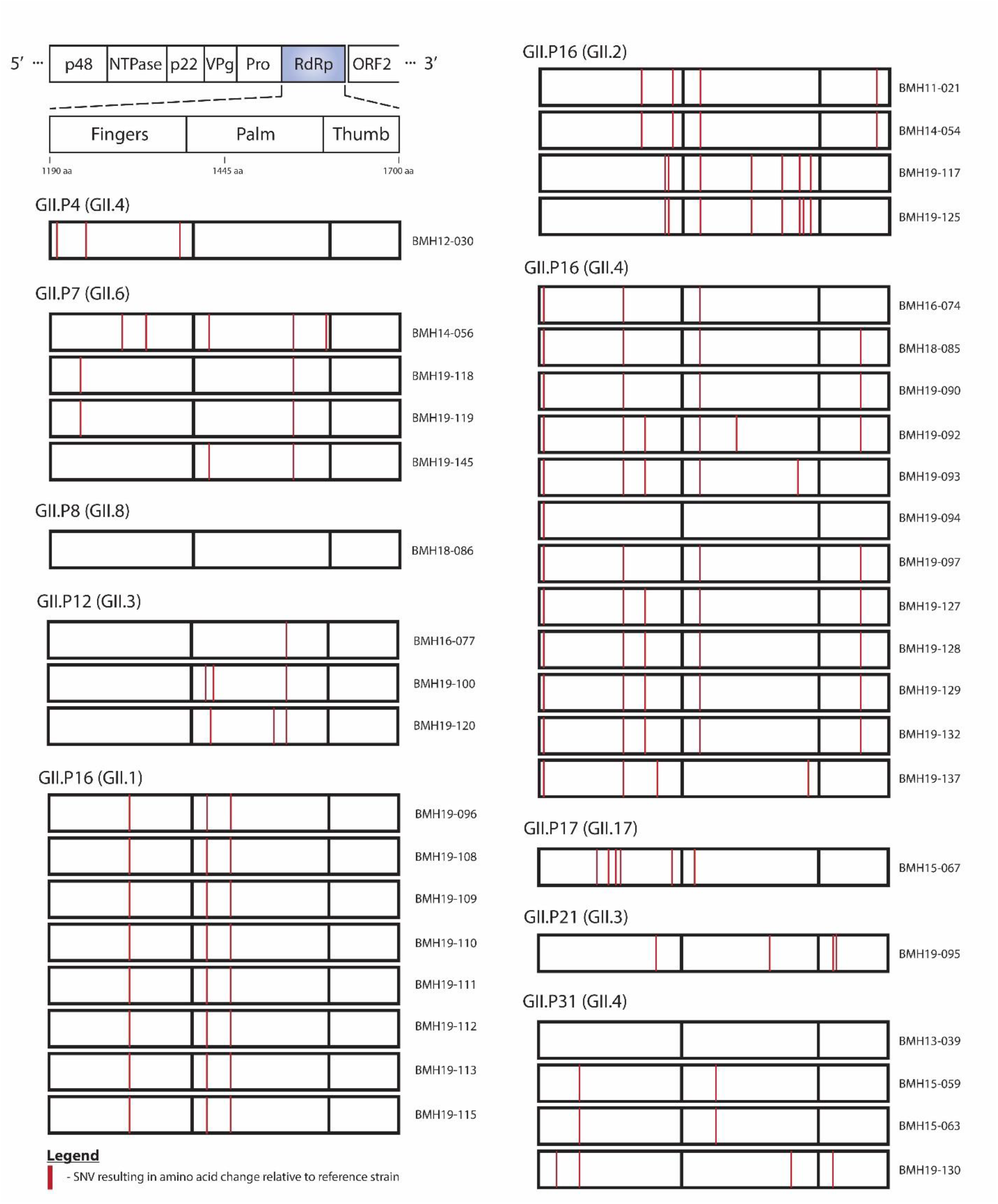
SNV analysis of norovirus RdRP (Polymerase) region. Norovirus *de novo* assemblies were assessed for variants relative to highly similar NCBI reference sequences. Schematic representation of the norovirus RdRP region is shown (upper left). SNVs that resulted in non-synonymous amino acid changes are shown (red lines) at corresponding amino acid positions in the Fingers, Palm, and Thumb domains for each isolate. Isolates are grouped by polymerase type. The GenBank reference Isolates (accession numbers in brackets) used were GII.P4-2011-USA (KC576912), GII.P7-2018-USA (MT731279), GII.P8-2018-China (MN996298), GII.P12-2015-USA (MT409884), GII.P16-2016-USA (MK753033), GII.P16-2011-USA (KJ407074), GII.P16-2017-USA (MK762635), GII.17-2016-China (KU757046), GII.P21-2015-UK (MH218671), and GII.P31-2015-Canada (KX158279).

### Recombination

Noroviruses are prone to recombine at the boundary of ORF1/ORF2 (Parra. 2019). To investigate recombination events, the 39 whole genome norovirus sequences were analyzed using RDP4 (Figure 6 and Supplementary Table 3). As shown in Figure 7a, BMH16-074 (GII.4 Sydney 2012[P16] was identified as being a recombinant strain. The major parent has the genotype GII.1[P16] (GenBank accession MK753033) and minor parent is GII.4 Sydney 2012[P31] (GenBank accession KX158279). Thus, a recombinant event at the ORF1/ORF2 boundary resulted in BMH16-074 obtaining the ORF2/3 region for GII.4 Sydney 2012. Additionally, the strains BMH18-085, BMH19-090, BMH19-092, BMH19-093, BMH19-094, BMH19-097, BMH19-127, BMH19-128, BMH19-129, BMH19-132 and BMH19-137 show evidence of this same recombination event (Supplementary Table 4).

**Figure 7.**
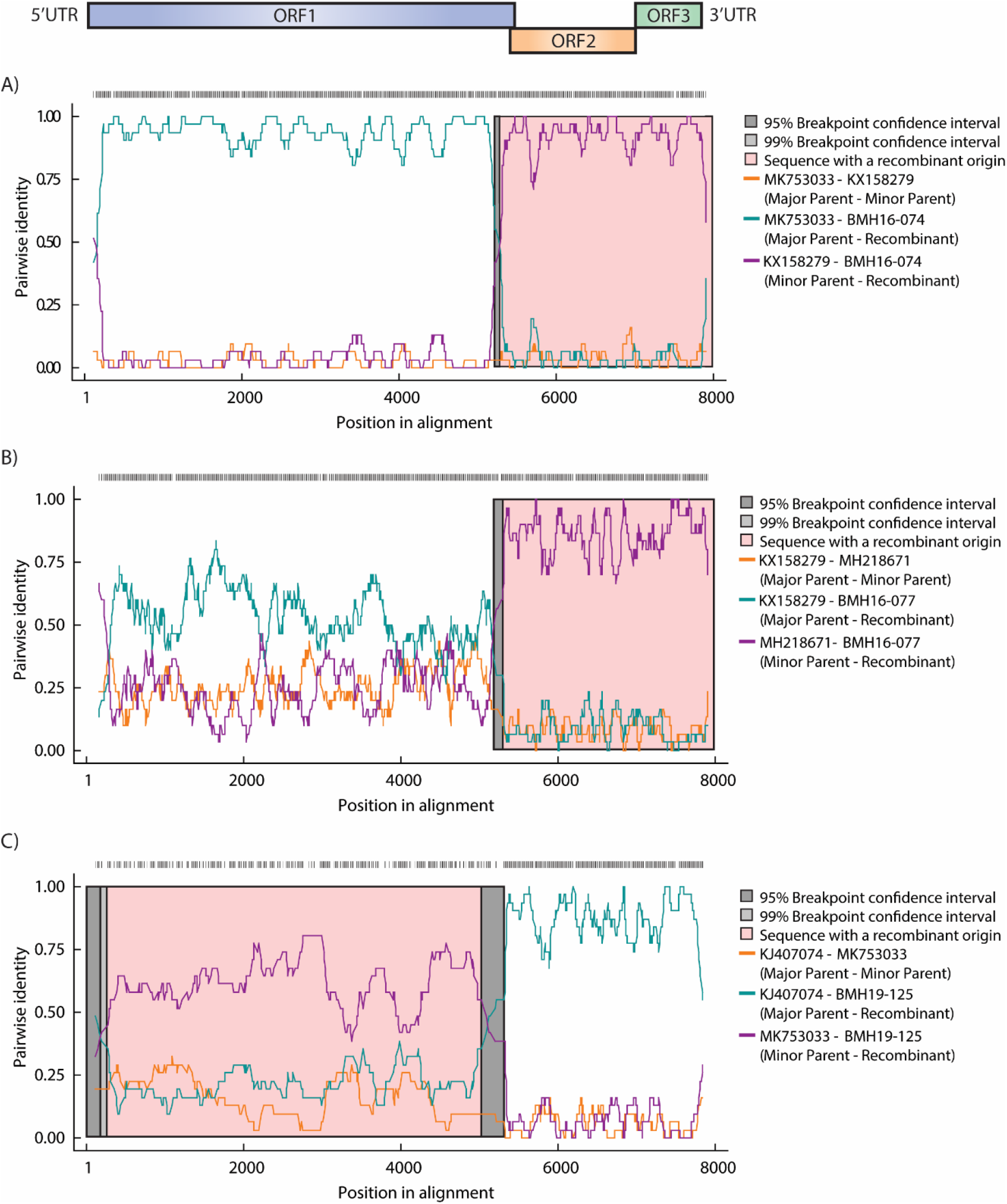
Recombination analysis of norovirus strains used in this study. Simplots were constructed using RDP4 (v4.101) by comparison of the complete *de novo* norovirus genomes to the reference strains using a slide window width of 200 bp and step size of 20 bp. (A) Percentage similarity of recombinant BMH16-074 to parental reference strains (Genbank accession MK753033 and KX158279). (B) Percentage similarity of recombinant BMH16-077 to parental reference strains (Genbank accession KX158279 and MH218671). (C) Percentage similarity of recombinant BMH19-125 to parental reference strains (Genbank accession KJ407074 and MK753033). Schematic representation of the norovirus genome is shown above the graph.

BMH16-077, genotype GII.3[P12], shows evidence of a major recombination event at the ORF1/ORF2 boundary (Figure 7b). GenBank reference strain MH218671, which has the genotype GII.3[P21], was identified as the minor parent, and reference strain KX158279 with genotype GII.4 Sydney 2012 [P31], the major parent. The recombination event resulted in BMH16-077 exchanging the GII.4 Sydney 2012 ORF2 region for the GII.3 sequence. BMH19-100 and BMH19-120 also show evidence of the same recombination event (Supplementary Table 4).

Evidence of a recombination event at the ORF1/ORF2 boundary is also observed for BMH19-125 (Figure 7c). For this isolate, BMH19-125, which has the genotype GII.2[P16], underwent a recombination event with the GenBank reference strain MK753033 (GII.1[P16]). The major parent identified was GenBank reference isolate KJ407074 (GII.2[P16]). This event resulted in the recombination of the ORF1 GII.P16 between BMH19-125 and GenBank strain MK753033. Evidence of this same recombination event is also observed in BMH19-117 (Supplementary Table 4).

## Discussion

The approach implemented by Parra and colleagues for full-length amplicon generation of GII samples has the potential to revolutionize the genome analysis of noroviruses. Herein, we applied this approach on 57 norovirus GII positive samples and obtained full-length amplicons for 39 samples (>67%). We performed Sanger sequencing to determine the genotype of the remaining samples. The failure of these samples to generate full-amplicons could be explained by the presence of RT-PCR inhibitors, as the viral titre for these samples are above the limit of the assay. For future applications, diluting the samples could potentially reduce the effect of RT-PCR inhibitors and aid in obtaining full-length amplicons (Nasheri, et al. 2020). We have also determined the lowest viral RNA load that would render full-genomic amplicons for four representative samples, and demonstrated that 170 to 346 genome copies would be enough for WGS using this technique. This is promising as the level of natural contamination for certain high-risk foods such as oysters is within or even higher than this range (Le Guyader, et al. 2009). Therefore, this approach has the potential to be employed for WGS analysis of naturally contaminated food products in the absence of an enrichment strategy.

Coverage analysis of representative norovirus genomes demonstrated the utility of combining long read Nanopore and short read Illumina sequencing data to obtain full-length *de novo* norovirus genomes. Using an approach that consisted only of Illumina sequencing, we often obtained incomplete genome assemblies. Indeed, decreased sequencing coverage at the 5’ and 3’ end of the norovirus sequence is observed for the Illumina data (Figure 3), resulting in partial assembly for ORF1, and less frequently, ORF3 (data not shown). By using a hybrid genome assembly approach, we were able to obtain highly accurate complete norovirus genomic sequences by first extracting full-length Nanopore sequence reads followed by error correction using a combination of both long and short read sequencing data. Differences in coverage depth along the genomes that are observed in some of the samples (Figure 3) are likely the result of incomplete cDNA synthesis (partial genome length amplicons), or the presence of subgenomic sequences.

In recent years, the emergence and spread of RdRp/capsid recombinant noroviruses have been reported around the world. In the United States, a new recombinant GII.4 Sydney emerged in 2015 (GII.4 Sydney[P16]) and replaced the 2012 variant (GII.4 Sydney[P31], formerly GII.Pe-GII.4 Sydney), which was the dominant strain for several years throughout the world (Barclay, et al. 2019). In Alberta, Canada, GII.4 Sydney[P16] was predominant in 2015–2016 and 2017–2018 (Hasing, et al. 2019), GII.2[P16] was predominant in 2016–2017 (Hasing, et al. 2019), and GII.12[P16] emerged in 2018–2019 and caused 10% of outbreaks and 17% of sporadic cases (Pabbaraju, et al. 2019). Herein, GII.4 Sydney[P16] made up 13 out of 30 (43.3%) samples since 2016 and thus was the predominant strain in 2016-2019, consistent with what has been reported before (Hasing, et al. 2019). Although, a combination of other genotypes such as GII.3[P12], GII.1[P16], and GII.6[P7] were co-circulating at this time, GII.4 isolates were only associated with GII.P16 (Supplementary Table 1). It was also interesting to detect the GII.17[P17] strain that caused several outbreaks in multiple countries during 2014 and 2015 (Matsushima, et al. 2019, van Beek, et al. 2018) from a sample that was isolated in 2015 (Supplementary Table 1).

Even though GII.P16 has recently re-emerged as the dominant polymerase type, the oldest isolate in this study BMH11-021 also has P16, but it does not cluster with the recent isolates, instead it shows homology to older P16 isolates from 2011 (Figure 4 A). On the other hand, more recent GII.4 isolates from 2019 still show high homology to the older isolates from 2013, 2014, and 2015. This observation, together with the fact that GII.4 has continued to be the dominant genotype for more than three decades, indicates that despite some sequence plasticity, the GII.4 capsid is evolutionarily conserved due to some fitness advantages. Our results also demonstrate that samples BMH19-108 to BMH19-115 resulting from the GII.1[P16] outbreak are highly homologous across all three ORFs (Figure 4).

The P2 subdomain of the VP1 protein interacts with potential norovirus carbohydrate receptors and contains 6 antigenic epitopes (Debbink, et al. 2012, Lindesmith, et al. 2013), therefore, it is not surprising to see that the majority of the non-synonymous SNVs are concentrated in this subdomain (Figure 5). The GII.4 Sydney[P16] isolates in this study, all contain the V377A change in the antigenic sites in the P2 subdomain (Supplementary Table 2) already described among GII.4 Sydney[P16] isolates in the U.S. (Cannon, et al. 2017), which is absent in GII.4 Sydney[P31] isolates. In addition, BMH19-090 contains the M333V change in the epitope C of the P2 subdomain that was reported in GII.4 Sydney[P16] isolates in multiple studies (Cannon, et al. 2017, Ruis, et al. 2020). It has been suggested that these substitutions influence viral fitness by altering antigenicity, increasing transmissibility, receptor binding, or particle stability (Ruis, et al. 2020). Outbreak samples BMH19-127 to BMH19-132 have a S470T within the P1 subdomain that has not been reported before. GII.2[P16] strains have been reported in the last decade, and the S24N, and V335I substitutions observed in the isolates in this study have been detected before (Tohma, et al. 2017). However, V256I, V319I, V373I, and V440I substitutions were only observed in the samples from this study (Supplementary Table 2), and they are considered biochemically conserved substitutions.

The re-emerging P16 isolated in this study were associated with three capsid genotypes; GII.4, GII.2, and GII.1 (Figure 6). The outbreak associated GII.P16 (GII.1) isolates all have the same three substitutions T1294S, S1401T, and V1446A (supplementary Table 3). However, the S1401T substitution in the Palm domain was also observed in GII.P16 (GII.2) and GII.P16 (GII.4) isolates. Three out of four sporadic P31 isolates in this study, which are associated with GII.4 Sydney capsid, had R1236K substitution in the Fingers domain. The GII.P12 isolates all have the V1521I substitution, and all the GII.P7 isolates have the I1519V substitution within the Palm domain, however, none of these substitutions falls within the conserved motifs or the active sites, and they are all considered biochemically conserved substitutions, thus their significance is not known.

In conclusion, using the full-length amplicons is an efficient and sensitive method for norovirus WGS analysis. Also, our results are consistent with other reports regarding the predominance of GII.4[P16] Sydney replacing the previous GII.4[P31] Sydney, indicating the enhanced fitness of the GII.4 Sydney capsid. Continued norovirus genomic surveillance will help in the understanding of norovirus evolutionary mechanisms and the identification of emerging variants, which ultimately aid in designing future norovirus vaccines and antivirals.

## Acknowledgements

The authors would like to thank Dr. Brent Dixon and Dr. Ana Pilar from the Bureau of Microbial Hazards for kindly reviewing the manuscript and providing insightful comments. This study is financially supported by the Bureau of Microbial Hazards, Health Canada.

